# The CST complex mediates a post-resection non-homologous end-joining repair pathway and promotes local deletions

**DOI:** 10.1101/2025.02.07.637039

**Authors:** Oana Ilioaia, Liébaut Dudragne, Clémentine Brocas, Léa Meneu, Romain Koszul, Karine Dubrana, Zhou Xu

## Abstract

Repair of a DNA double-strand break (DSB) by non-homologous end-joining (NHEJ) generally leaves an intact or minimally modified DNA sequence. Resection initiation exposes single-stranded DNA and directs repair towards homology-dependent pathways and away from NHEJ. Therefore, NHEJ is not thought to be an available repair pathway once the DSB is resected. Here, we report that the Cdc13/Stn1/Ten1 (CST) complex, well characterized for its telomere-associated functions, acts after resection initiation to mediate a backup NHEJ repair. We found a CST-specific mutation signature after DSB repair, characterized by deletions of 5-85 bp, mostly dependent on NHEJ. In contrast, NHEJ-mediated small deletions of 1-4 bp and insertions are not affected in CST mutants. The interaction between CST and Polα-primase is critical for these intermediate size deletions, suggesting a role for fill-in synthesis. Consistently, in *stn1Δ* and Polα-primase mutant deficient for interaction with CST, resection is increased, leading to larger deletions of several kilobases mediated by microhomologies. Collectively, these results depict a more complex picture of repair pathway choice where CST allows a post-resection NHEJ repair, promoting local deletions but guarding against much larger and potentially more deleterious deletions and rearrangements.

## Introduction

DNA damage, particularly double-strand breaks (DSB), poses a major threat to genome integrity. To deal with this threat, the DNA damage response (DDR) is activated to ensure appropriate repair (Finn et al., 2012; Harrison and Haber, 2006). Following a DSB, an important step of the DDR is the processing of the break by resection of the 5′ extremities, which controls the repair pathway choice between two main repair mechanisms (Symington and Gautier, 2011): homologous recombination (HR), which requires single-stranded DNA (ssDNA) exposure for homology search and strand annealing, and non-homologous end-joining (NHEJ), which directly ligates the DSB ends without extensive processing. Two other mechanisms distinct from HR and relying on sequence homology can join DSB ends: single-strand annealing (SSA) and microhomology-mediated end-joining (MMEJ). Both require resection to expose the homologous sequences and lead to deletions, but differ in several aspects of their molecular mechanisms and in their genetic requirements.

### CST, a major player in telomere protection and maintenance

In contrast, telomeres, the natural extremities of eukaryotic linear chromosomes, resemble one side of a DSB but do not trigger a DDR, which would lead to inappropriate repair and genome instability (Wellinger and Zakian, 2012). They are thus protected by proteins, bound to the double-stranded and the single-stranded parts of the telomere, that inhibit the DDR. In *Saccharomyces cerevisiae*, the CST complex, composed of Cdc13/Stn1/Ten1, binds and protects the single-stranded TG_1–3_ telomeric repeats, prevents Exo1 from resecting the 5’ strand, and recruits telomerase by direct interaction with Est1 (Bianchi et al., 2004; Churikov et al., 2013; Gao et al., 2007; Garvik et al., 1995; Grandin et al., 2001; Grandin et al., 1997; Nugent et al., 1996; Pennock et al., 2001; Qi and Zakian, 2000). In addition, CST recruits Polα-primase for lagging strand synthesis to resynthesize the double-stranded DNA (dsDNA) after replication or telomerase activity (Grossi et al., 2004; Qi and Zakian, 2000). Collectively, these roles prevent telomere loss and the associated chromosomal instability to preserve genome integrity.

### Functions of CST in DNA repair

While CST’s role at telomeres is well-established, its extra-telomeric functions are less well understood. The human CST complex (CTC1/STN1/TEN1) has been implicated in recovery and genome stability after replication stress, by promoting new origin firing or stimulating POLα for replication restart (Stewart et al., 2018). The CST complex was also shown to stabilize and protect stalled replication forks (Chastain et al., 2016; Jaiswal et al., 2023; Lyu et al., 2021). Recently, the CST complex together with Polα-primase has been involved in DSB processing as an effector of the 53BP1-Rif1-Shieldin axis, which controls resection and repair pathway choice, by performing fill-in synthesis on 3’ overhangs and facilitating NHEJ repair (Barazas et al., 2018; King et al., 2024; Mirman et al., 2018; Mirman et al., 2022; Schimmel et al., 2021). Additionally, CST was reported to promote DNA repair and survival in response to oxidative damage (Hara et al., 2023; Wysong et al., 2024).

In budding yeast, CST’s interaction with Polα-primase was suggested to regulate transcription during replication (Calvo et al., 2019). Additionally, Cdc13 localizes at DSBs in a Mre11- and Rad51-dependent manner, suggesting it associates with resection-mediated ssDNA (Bianchi et al., 2004; Horigome et al., 2014; Oza et al., 2009; Zhang and Durocher, 2010). Cdc13 can then recruit telomerase, promoting telomere addition even in absence of telomeric sequences, albeit with less efficiency (Bianchi et al., 2004; Diede and Gottschling, 1999, 2001; Kramer and Haber, 1993).

However, whether the CST complex contributes to DSB repair in budding yeast is not known. Direct genetic approaches to address this question are hindered by the essentiality of CST’s telomeric functions. In this work, we set up an experimental system with an inducible Cas9 DSB in a yeast strain with a single circular chromosome devoid of telomeres, in which the CST is no longer essential for survival. Through a comprehensive analysis of mutation signature after DSB repair, we demonstrate that the CST complex, and prominently Stn1, contributes substantially to NHEJ-mediated repair of the DSB through its interaction with Polα-primase. More specifically, we show that CST acts after resection initiation and is critical for intermediate size deletions (5-85 bp) formed through NHEJ. Altogether, we reveal an important role of the CST complex in DSB repair, as the mediator of a back-up NHEJ pathway that acts after resection.

## Results

### Inducible Cas9 DSB in a single circular chromosome strain

To uncouple the role of the CST complex at a DSB from its telomeric functions, we generated a yeast strain devoid of telomeres (Figure 1A). We took advantage of a strain where all 16 chromosomes are fused into a single linear one (SY14) (Shao et al., 2018) to build a strain with a single circular chromosome. To do so, we induced two simultaneous Cas9 cuts at the two remaining subtelomeres and recombined them with a chimeric oligonucleotide template homologous to both sequences. The successful chromosome fusion and the absence of telomeres were confirmed by Southern blot (Figures S1A and S1B). Chromosome conformation capture analysis by Hi-C confirmed the circular nature of the single chromosome and showed no other significant structural difference between the single linear and single circular chromosomes (Figure S1C).

**Figure 1.**
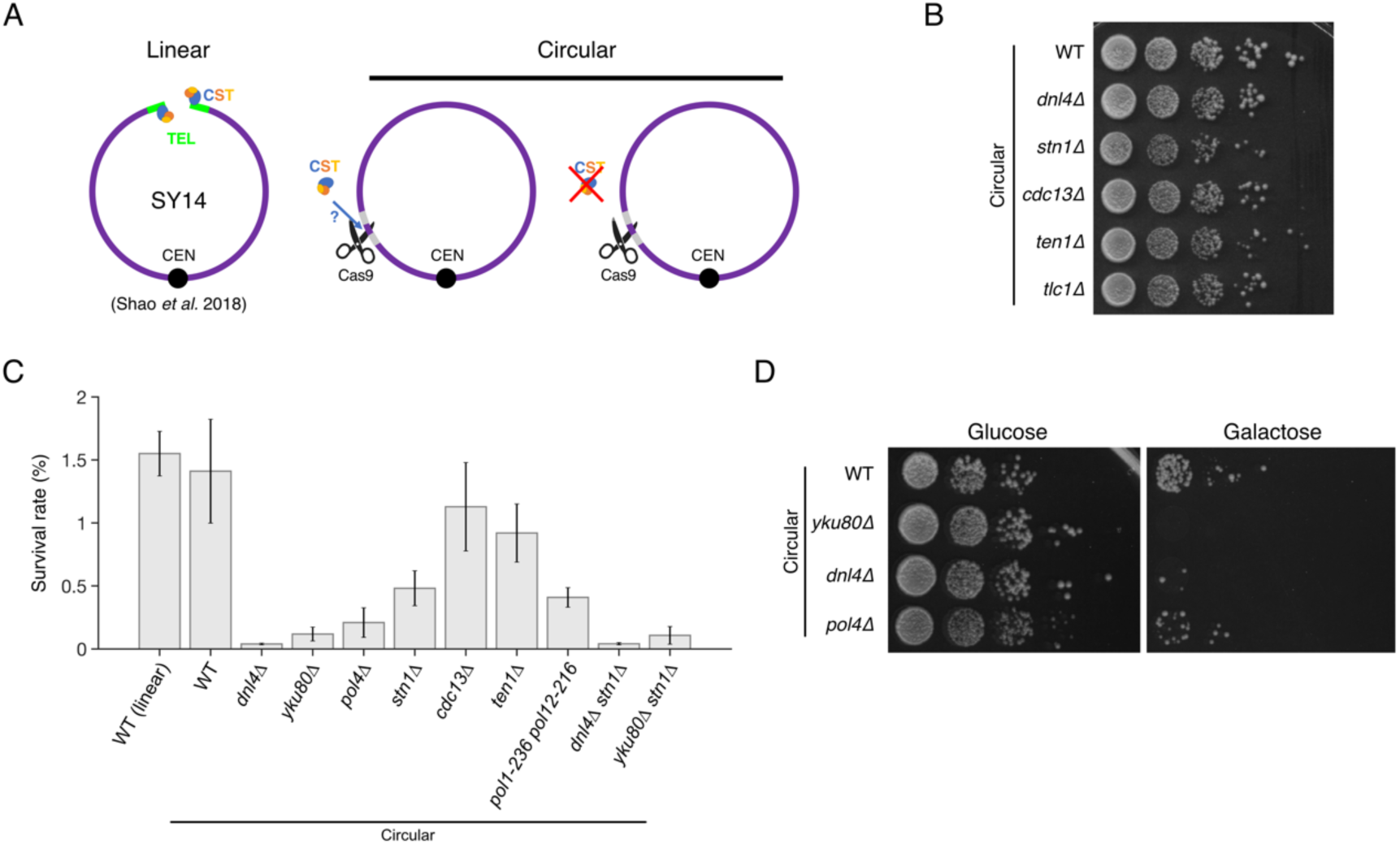
A yeast strain with a single circular chromosome and an inducible DSB as an experimental system. (A) Schematic representation of the yeast strain with a single linear chromosome (SY14, left) (Shao et al., 2018) and the one built in this work with a single circular chromosome and an inducible Cas9 DSB (middle and right), allowing for viable deletion of any CST subunit gene. (B) Spot assay with the indicated strains showing cell viability on rich YPD media. (C) Survival rate after DSB induction for the indicated strains. Mean and average are shown for n = 3-6 independent experiments. (D) Spot assay showing cell survival and growth with (galactose-containing plate, right) or without (glucose-containing plate, left) Cas9 induction for the indicated strains.

As also shown previously by others (Shao et al., 2019; Wu et al., 2020), a single circular chromosome strain is viable without telomerase, confirming that telomere maintenance was no longer needed (Figure 1B). Similarly, telomere protection was no longer essential since the deletion of any of the CST genes did not impair cell survival (Figure 1B), as observed previously (Wu et al., 2020).

To investigate DSB repair, we targeted, in the single circular chromosome strain, a unique site in the 5’ UTR of the *LYS2* gene using a plasmid expressing a specific guide RNA and a galactose-inducible Cas9 (Figure 1A). DSB induction was triggered by plating cells on galactose-containing media and survival was quantified by spot assays and colony-forming units relative to control cells plated on glucose. Since Cas9 is continuously expressed in galactose-containing media, repair of this DSB requires a mutational event to prevent further cutting by Cas9 and allow cell survival. Following DSB induction, survival of the wild-type (WT) circular chromosome strain (mean ± SD = 1.41 ± 0.82 %) was similar to the survival of the WT linear chromosome strain (1.55 ± 0.35 %) (Figures 1C and 1D), indicating that chromosome circularization did not significantly alter DNA repair efficiency. We thus used the circular chromosome strain together with this single Cas9-inducible DSB as an experimental system to investigate the specific roles of the CST complex in DSB repair.

### Repair of the Cas9 DSB occurs through NHEJ and MMEJ

We then characterized the mechanisms underlying repair in our experimental system. Survival after DSB induction primarily depended on the NHEJ pathway components ligase 4 (encoded by *DNL4*) and Yku80, as expected (Figures 1C and 1D). Survival was also largely dependent on Pol4, which is required for small fill-ins around the break and contributes significantly to the inaccurate NHEJ pathway (Tseng and Tomkinson, 2002; Wilson and Lieber, 1999).

In mammalian cells, repair of Cas9 DSBs has also been reported to produce large deletions (LDs) ranging from ∼100 bp to several kb (Kosicki et al., 2018; Owens et al., 2019). To evaluate the frequency of this outcome in our experimental system, we analyzed the colonies surviving DSB induction by multiplex PCR with one pair of primers flanking the DSB (amplicon size of 176 bp) and another one in an unrelated region of the genome to control for PCR efficiency, expecting that LDs would lead to unproductive PCRs. A PCR product around the cut site was amplified in 114 out of 115 (99.1%) WT colonies tested (Figure S2A). Limited variations of the size of the amplicon were observed and were consistent with NHEJ or MMEJ repair leading to local insertions and deletions. We further investigated the single unproductive PCR by PCR mapping and junction sequencing and found that, instead of a LD, it corresponded to an inversion of a 127-bp sequence close to the cut site associated with limited deletions at the boundaries (Figure S2B). Thus, no LD was found among 115 survivor colonies tested.

We then investigated the mutational signature at the Cas9 DSB site by high-throughput sequencing of a 231-bp amplicon across the DSB site from thousands of survivors, using a method developed by the Tijsterman lab (Schimmel et al., 2021; van Schendel et al., 2022). Such an approach has proved extremely powerful at characterizing the spectrum of DSB repair outcomes and dissecting their genetic determinants (Hussmann et al., 2021; Schimmel et al., 2021). Analysis of the repaired sequences revealed that WT cells with a circular chromosome mostly exhibited a mixture of deletions and insertions, with minor fractions of other types of mutations (deletion with insert, templated insertion and single-nucleotide variant (SNV)), similar to the linear chromosome strain (Figures 2A and S2C). In contrast to WT, NHEJ-deficient mutants (*dnl4Δ*, *yku80Δ* and *pol4Δ*) showed a dramatic disruption of the mutational signature, with a strong decrease in insertions and deletions in survivors, confirming repair by inaccurate NHEJ in WT (Figures 2A and S2B).

**Figure 2.**
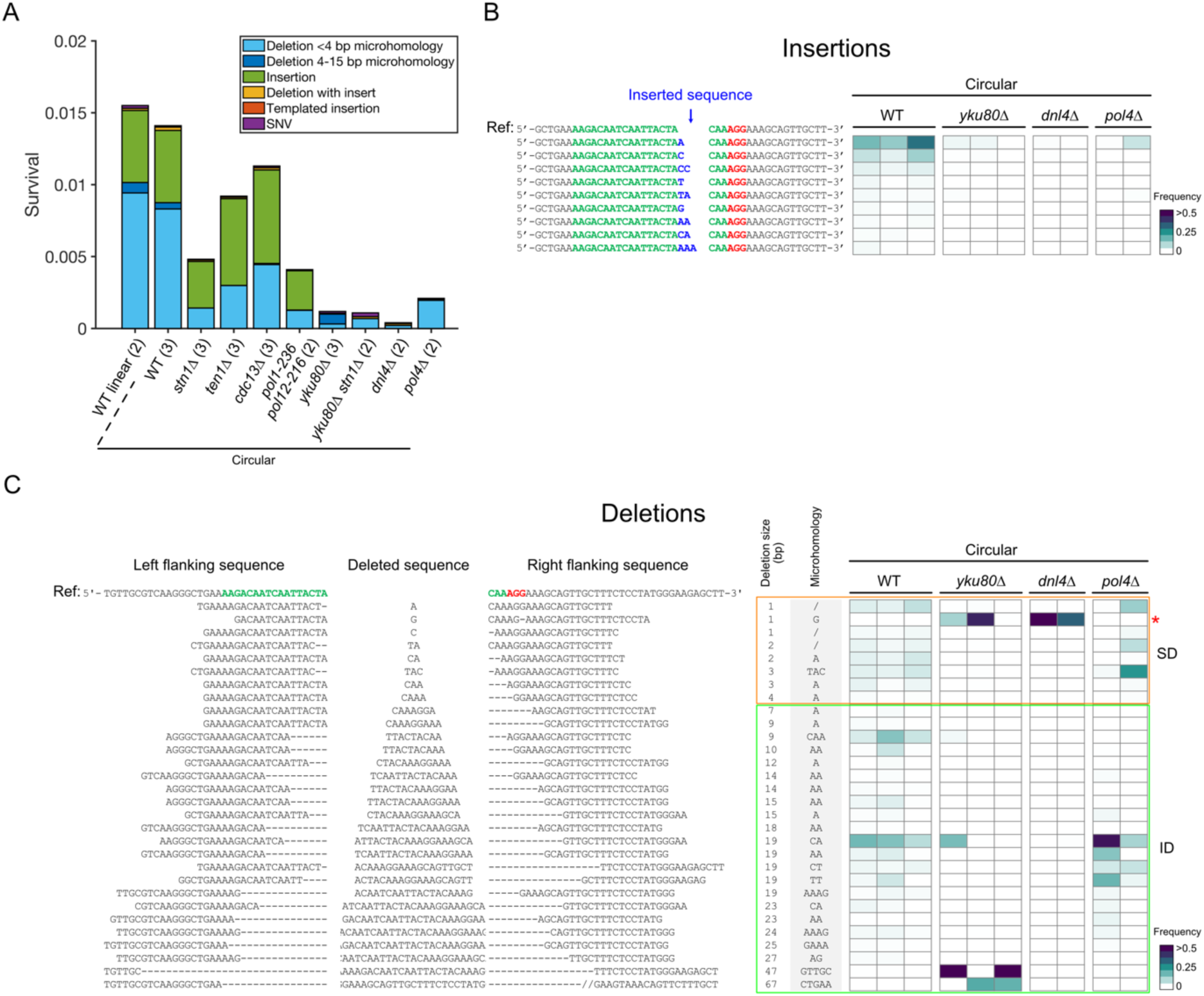
NHEJ and MMEJ mediate repair of the Cas9 DSB. (A) Deep sequencing analysis of mutation signature after DSB repair in the indicated strains, with fraction of different mutation types shown as stacked bars, normalized by survival rate. Between ∼200 and ∼3000 colonies surviving Cas9 DSB induction were collected in each experiment. Multiplex high-throughput Illumina sequencing of the amplicon around the DSB was performed. Analysis and clustering were done using SIQ (Schimmel et al., 2021; van Schendel et al., 2022). Parentheses show the number of independent experiments. (B) Heatmap of the frequency of each repair outcome with an insertion for individual experiments (in columns) with the indicated strains. The inserted sequences are shown in blue, the sequence targeted by the guide RNA in green and the PAM sequence in red. The insertions were ordered according to their increasing average frequency in the wild-type circular strain. Only insertions that appear with an average frequency of >0.005 are shown. The color bar shows the frequency scale. See Supplemental Data 1 and 2 for the unfiltered data. (C) Heatmap of the frequency of each repair outcome with a deletion for individual experiments (in columns) with the indicated strains. The deleted (cropped for size constraints) and flanking sequences are indicated. The deletions are ordered according to their increasing sizes. The “Microhomology” box shows whether microhomologies were found at the boundaries of the deletion, with the sequence indicated. Only deletions that appear with a frequency of > 0.01 in at least one experiment are shown. Red asterisks: deletion of one G located 4-5 bp away from the cut site, likely not due to the repair of the DSB. The color bar shows the frequency scale. See Supplemental Data 1 and 2 for the unfiltered data.

Upon closer examination of the sequences, we identified three predominant types of mutations in the WT circular strain at the repaired cut site: (i) insertion of a single base, (ii) deletion of 1-4 bases and (iii) deletion of 5-85 bases (Figures 2B and 2C).

We observed high-frequency insertions of a single nucleotide adjacent to an existing identical nucleotide at the Cas9 cut site (A in 19.5 ± 9.8 % and C in 7.0 ± 3.8 %) (Figure 2B). As proposed by Lemos and colleagues (Lemos et al., 2018), these insertions may be caused by the occasional non-blunt cutting by Cas9, leaving a 5’ overhang of one nucleotide, which would subsequently be filled by Pol4 and then ligated. In line with this hypothesis, the *pol4Δ* mutant showed a much lower frequency of these one-base insertion events (A: 3.3 ± 4.2 %; C: 0.026 ± 0.016 %) (Figure 2B). These insertions were also dependent on Yku80 and Dnl4, confirming that they resulted from NHEJ repair.

In WT cells, short deletion (SD) repair events (1-4 bp) occurred near the cut site and showed few (< 4 bp) or no microhomology at their boundaries, consistent with NHEJ-associated deletions. This was confirmed by their near-complete absence in *yku80Δ* and *dnl4Δ* mutants, with one notable exception: the deletion of a G from the PAM sequence, which occurred at very low frequency in WT cells (Figure 2C, red star). The position of this mutation, 4-5 bp away from the cut site, suggested that it was unrelated to the DSB itself and likely arose spontaneously before DSB induction in a NHEJ-independent manner. It was then selected because it impaired Cas9 cleavage, and became more prominent in *yku80Δ* and *dnl4Δ* mutants due to their low survival rates.

Intermediate size deletions (ID) ranging from 5 to 85 bp were generally bidirectional and frequently removed the PAM and a substantial portion of the guide sequence, with some deletions being particularly favored (*e.g.* deletion of 9 bp using CAA as microhomology and deletion of 19 bp using CA as microhomology), suggesting sequence-specific preference (Figure 2C). Among these IDs, 3.1% exhibited 4 or more bp of tandem microhomologies suggestive of MMEJ repair. MMEJ usage was supported by their enrichment in the *yku80Δ* mutant (57.9%) and dependence on ligase 4 and Pol4 (Figures 2C and S2C) (Lee and Lee, 2007; Ma et al., 2003). Surprisingly, the remaining IDs (40.2%) were not associated with significant microhomology (< 4 bp) and required Yku80 and Dnl4, indicating that they arose from NHEJ repair (Figure 2C). The deletion size of 5-85 bp suggested that repair occurred after some processing of the break, likely by short range resection.

Overall, the Cas9 DSB led to both NHEJ- and MMEJ-dependent mutations, including insertions and both SDs (1-4 bp deletions) and IDs (5-85 bp deletions). This experimental setup allowed us to investigate CST’s contribution to these repair events.

### CST contributes to inaccurate NHEJ repair

Since Cdc13 can recruit telomerase and could thus allow the propagation of a relinearized chromosome as a viable repair outcome, we first wondered whether telomere healing contributed to survival after DSB induction in our system. However, the telomerase-negative *tlc1Δ* mutant did not affect survival (Figure S3A), which, added to the observation that the only unproductive PCR around the DSB in WT survivors could be assigned to an inversion (Figures S2A and S2B), indicated that telomere healing was not a significant survival pathway in this experimental setting.

To investigate the implication of the CST complex in DSB repair, we deleted each of the 3 subunit genes and found that survival after DSB induction decreased, with the *stn1Δ* mutant showing the strongest effect (Figure 1C and 3A). We found a similar result when the DSB was induced at another locus, *i.e.* the 5’ UTR of *URA3* (Figure S3B). Combining *STN1* deletion with *dnl4Δ* or *yku80Δ* did not further decrease survival, indicating that Stn1’s contribution to repair was NHEJ-dependent (Figures 1C and 3A). CST is thus an important contributor to inaccurate NHEJ.

**Figure 3.**
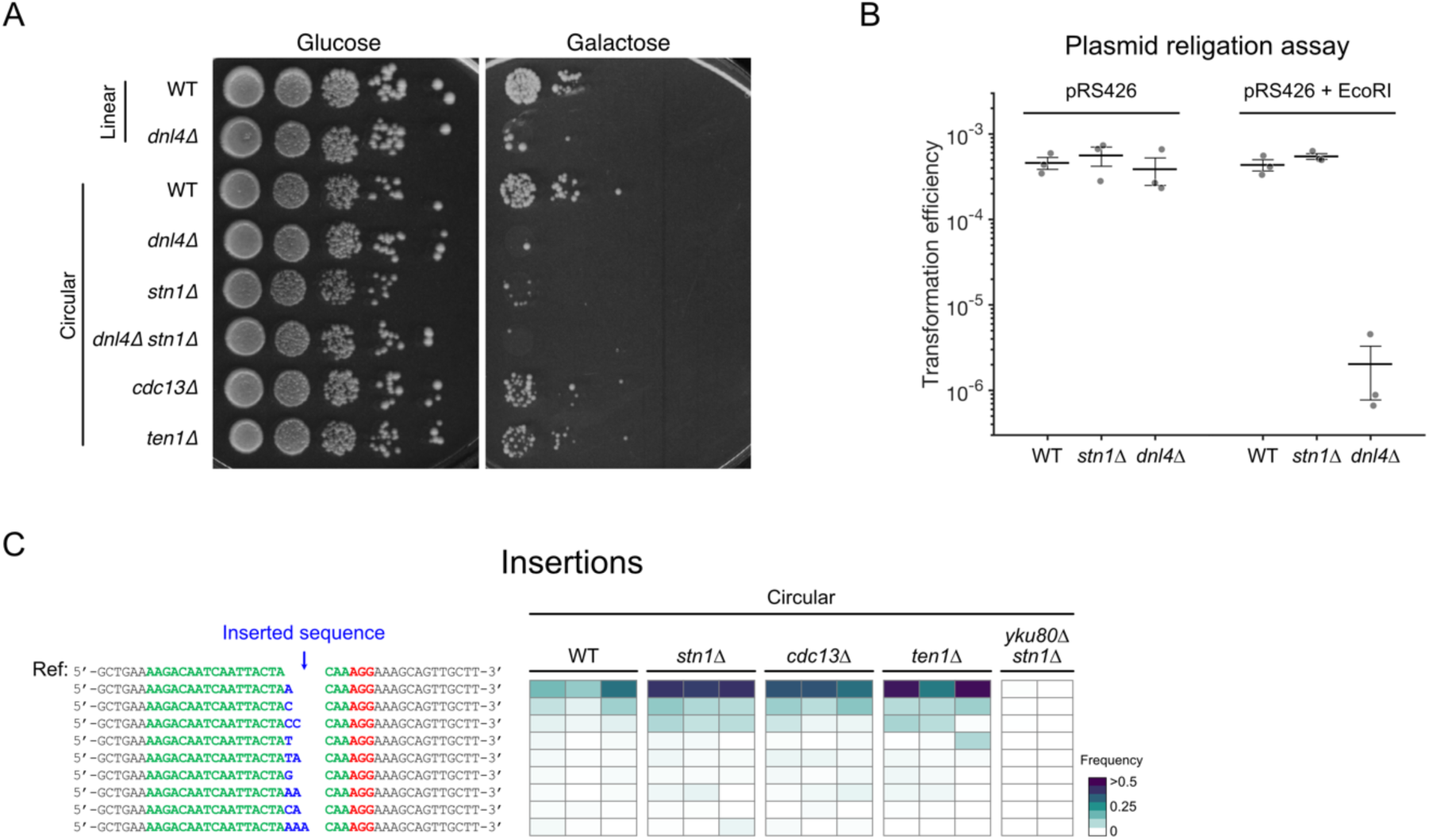
CST contributes to inaccurate repair of the DSB. (A) Spot assay as in Figure 1D with the indicated strains, including CST deletion mutants. (B) Plasmid religation assay. Transformation efficiency corresponds to the number of colonies formed on the selective plate without uracil normalized by the number of plated cells, as assessed on a non-selective YPD plate. Each dot represents an independent transformation experiment. Error bars show the standard error of the mean and the middle bar represents the mean. The strains were transformed either with the circular plasmid pRS426 or with the same plasmid linearized by EcoRI digestion (“pRS426 + EcoRI”). (C) Heatmap of the frequency of each repair outcome with an insertion for individual experiments (in columns) with the indicated strains as in Figure 2B. See Supplemental Data 1 and 2 for the unfiltered data.

To test whether CST also affects error-free NHEJ, we performed a plasmid religation assay. Circular plasmid transformation efficiency was similar in WT, *stn1Δ* and in *dnl4Δ* (Figure 3B). However, upon linearization, transformation efficiency dropped by ∼200 folds in *dnl4Δ*, as expected, but not in *stn1Δ*, indicating that Stn1 did not play a significant role in error-free NHEJ.

Thus, the CST complex does not affect error-free NHEJ but plays a telomerase-independent role in DSB repair through inaccurate NHEJ.

### CST does not affect NHEJ-mediated insertions

To precisely dissect the specific NHEJ pathway in which CST plays a role, we first asked whether CST affected NHEJ-mediated insertions. Examination of the mutation signature from high throughput sequencing in CST deletion mutants revealed that the mutation signature at the repaired cut site differed from both WT and NHEJ mutants (Figures 2A and S2C). The overall rate of insertions adjusted for survival was comparable to that of WT or slightly decreased in *stn1Δ* (Figure 2A). Detailed analysis confirmed that CST mutants did not alter the distribution of the most frequent insertions compared to WT, with single nucleotide insertions of A or C remaining the most frequent and being still Yku80- dependent (Figure 3C). This finding was corroborated at a different DSB locus (5’ UTR of *URA3*) where the distribution of insertions remained similar between WT and *stn1Δ* (Figure S3C). We thus conclude that CST does not influence NHEJ-mediated small insertions.

### CST is specifically required for intermediate size deletions

We next investigated CST’s contribution to all 3 deletion size ranges we defined: LDs, IDs and SDs. Using the multiplex PCR assay, we found that 11 out of 88 (12.5%) *stn1Δ* colonies led to unproductive PCR results (Figures 4A, S4A and S4B). Sequencing a random subset of 8 of these clones revealed large deletions of 924 (n = 1), 7551 (n = 5), 8777 (n = 1) and 8778 bp (n = 1), involving microhomologies of 5, 22, 11 and 11 bp, respectively (Figure S4B), likely resulting from MMEJ repair. Thus, Stn1’s activity limits MMEJ-mediated LDs.

**Figure 4.**
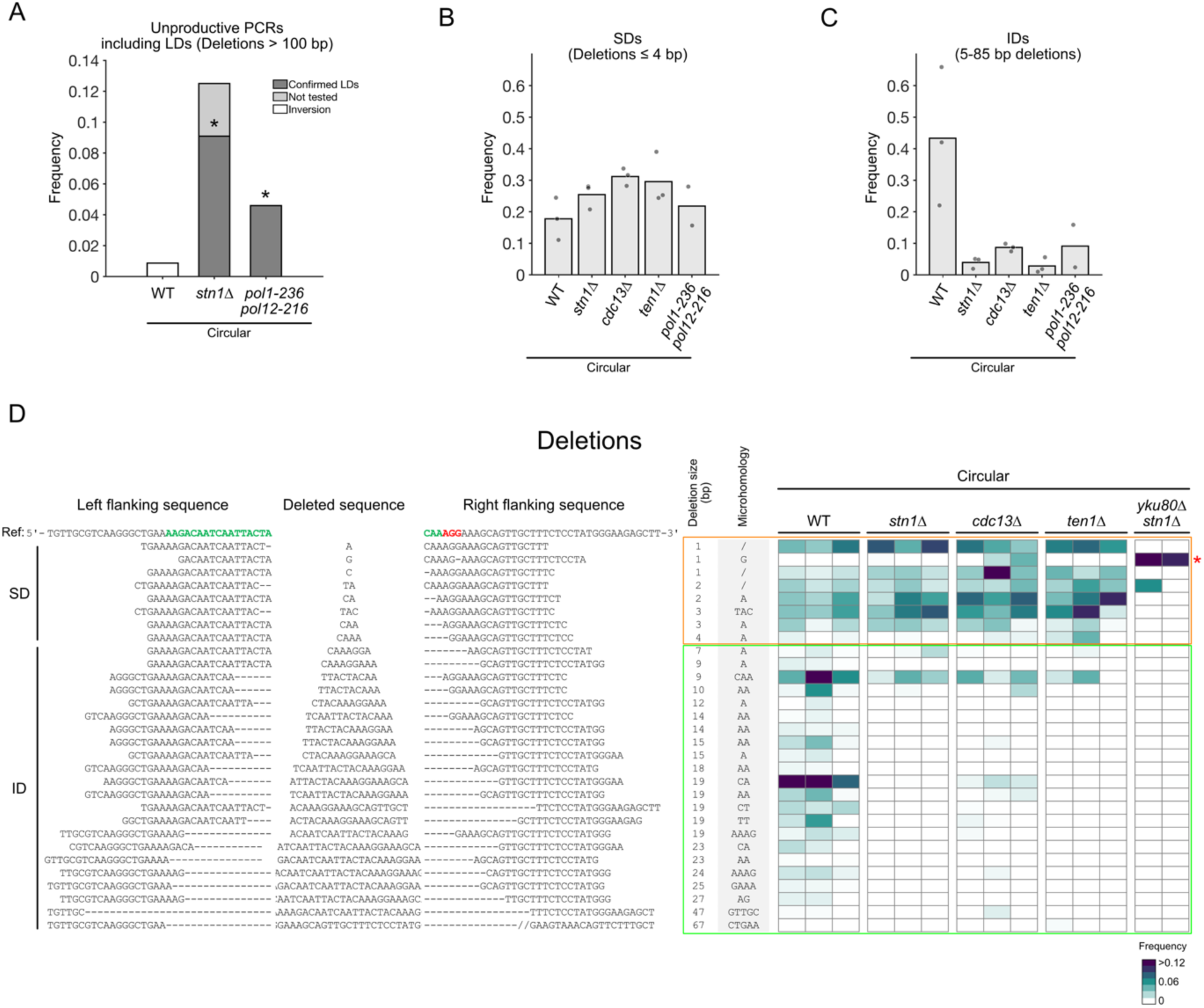
CST specifically affects intermediate size deletions. (A) Frequency of unproductive PCRs around the cut site in the indicated strains. Sequencing of the junction for the tested colonies revealed an inversion in the WT (see Figure S2B) and LDs in *stn1Δ* and *pol1-236 pol12-216* (see Figures S4A and S5). * indicate p-value < 0.05 compared to WT using two-tailed Fisher’s exact test on LD frequencies. (B) Frequency of SDs, *i.e.* deletions ≤ 4 bp, for the indicated strains. Each dot represents an independent experiment. (C) Frequency of IDs, *i.e.* 5-85 bp deletions, for the indicated strains. (D) Heatmap of the frequency of each repair outcome with a deletion for individual experiments (in columns) with the indicated strains, represented as in Figure 2C, except for the color bar which uses a different scaling. See Supplemental Data 1 and 2 for the unfiltered data.

In stark contrast to these LDs, the deletions captured by high throughput sequencing were significantly decreased overall in the 3 CST mutants, although to different extents, with *stn1Δ* again showing the strongest effect (Figure 2A). However, not all deletions required the CST complex. NHEJ-mediated SDs were not decreased in CST mutants, whereas IDs decreased by 5-10 folds in CST mutants compared to WT (Figures 4B, 4C and 4D). This finding was confirmed at another DSB locus, where IDs were also significantly decreased in *stn1Δ* mutants (Figures S4C and S4D).

Among IDs, those with larger microhomologies (4-15 bp) were drastically reduced in CST mutants compared to wild-type, suggesting decreased MMEJ repair (Figures 2A and S2B). This was further confirmed by the disappearance of the microhomology-associated IDs enriched in *yku80Δ* upon deletion of *STN1* (Figure S2B; compare also *yku80Δ* in Figure 2C with *yku80Δ stn1Δ* in Figure 4D). Likewise, deletions with few or no homologies (< 4 bp) were reduced both in frequency and when adjusted for survival (Figures 2A and S2B), demonstrating that intermediate size NHEJ-mediated deletions require CST.

Altogether, these data indicate that CST’s activity leads to distinct outcomes depending on the size of the deletions: no effect on SDs, stimulation of IDs and inhibition of LDs. Since CST promotes IDs regardless of the mechanism, NHEJ or MMEJ, used to complete repair, one hypothesis would be that CST acts upstream of repair per se, likely by limiting the formation of ssDNA, the extent of which would define deletion size and thus the formation of ID or LD. Considering CST’s interaction with Polα- primase and its role in post-replication fill-in synthesis at resected telomeres, we wondered whether CST functions after DSB resection, to recruit Polα-primase and regulate ssDNA.

### The CST complex limits resection by recruiting Polα-primase for fill-in synthesis

To test the relationship between CST and resection, we first asked whether limiting ssDNA formation could compensate for *stn1Δ*’s survival defect after DSB. We thus mutated the first step of resection dependent on the MRX complex and Sae2, either by deleting *SAE2* or by generating the nuclease-dead mutant *mre11-H125N*. As previously observed, survival by NHEJ repair significantly increased in these mutants (Figure 5A) (Deng et al., 2014; Lee and Lee, 2007). Deletion of *STN1* in these mutants did not decrease survival (Figure 5A), indicating that Stn1 is not required when resection is already limited. These results also suggest that CST acts downstream of resection in DSB repair.

**Figure 5.**
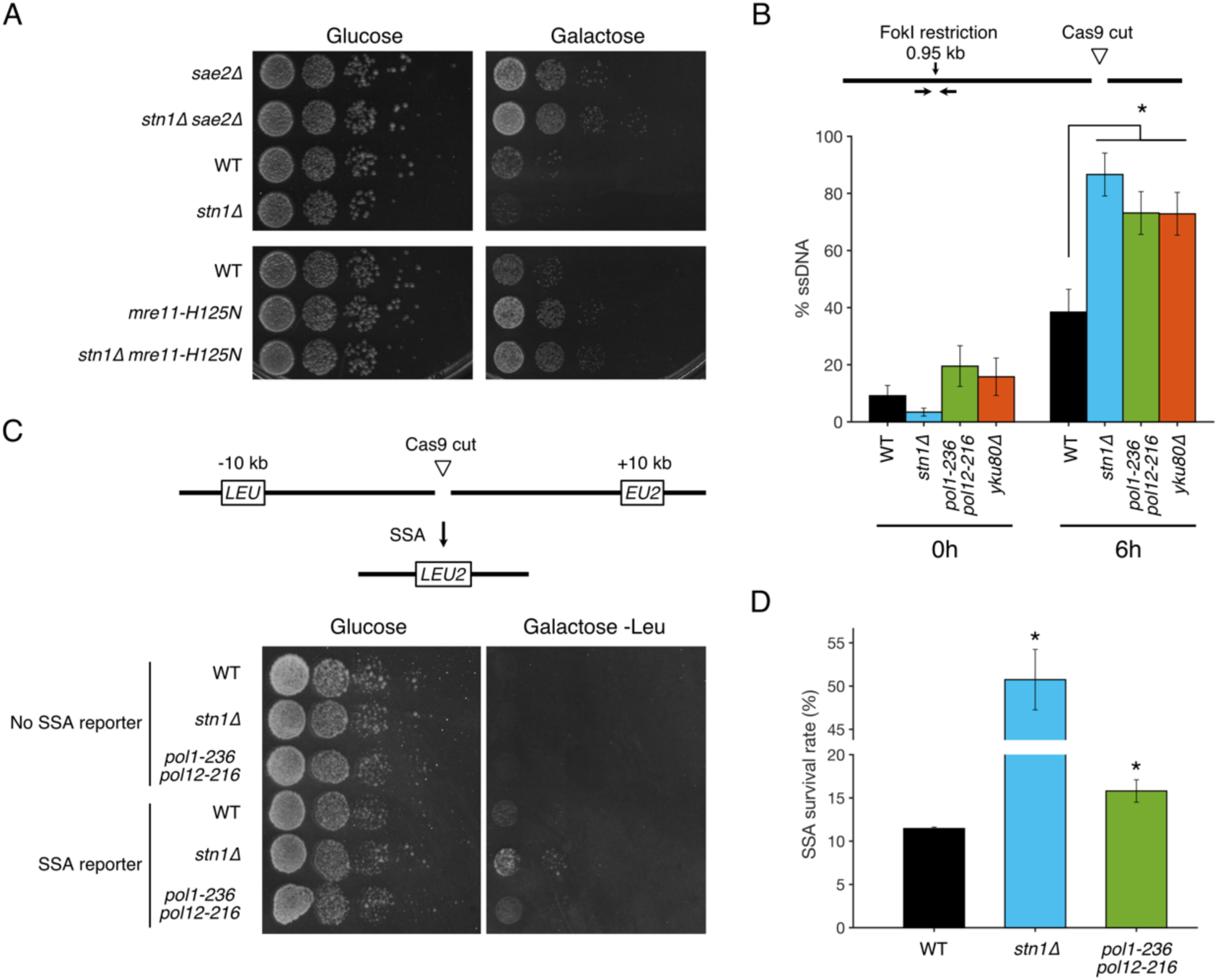
The CST complex limits resection through its interaction with Polα-primase. (A) Spot assay with the indicated strains showing cell survival and growth with (galactose-containing plate, right) or without (glucose-containing plate, left) Cas9 induction. (B) Quantification of ssDNA in the indicated strains (n = 5 independent experiments for WT and n = 3 for the others) at 0h and 6h after DSB induction, by qPCR after FokI digestion of the locus 0.95 kb away from the DSB, following the method described in (Zierhut and Diffley, 2008) and using within timepoint normalization. * indicates p-values <0.05 compared to WT using a two-tailed Student’s t-test. (C) SSA assay using reconstitution of *LEU2* as a readout by plating on media without leucine, with (right) or without (left) Cas9 induction. (D) Quantification of SSA survival rate (n = 3 independent experiments). * indicate p-values <0.05 compared to WT using a two-tailed Student’s t-test.

To determine whether the recruitment of Polα-primase by CST would affect the extent of ssDNA, we introduced point mutations in *POL1* (*pol1-236*: D236N) and *POL12* (*pol12-216*: G325D), which specifically disrupt the interaction between Polα and CST (Grossi et al., 2004; Puglisi et al., 2008; Qi and Zakian, 2000; Sun et al., 2011), and directly measured ssDNA accumulation 1 kb from the cut site by a restriction digest/qPCR method (Zierhut and Diffley, 2008). As a positive control, the *yku80Δ* mutant showed more ssDNA than WT after DSB induction, consistent with increased resection (Figure 5B) (Clerici et al., 2008; Lee et al., 1998). In *stn1Δ* and *pol1-236 pol12-216,* this assay also measured higher levels of ssDNA compared to WT (Figure 5B). Furthermore, using a SSA reporter, in which 82 bp of homology have been inserted at ∼10 kb on both side of the DSB and would allow the reconstitution of a functional *LEU2* gene after repair, we observed a ∼5-fold increased survival on plates lacking leucine in *stn1Δ* strain and a more modest but statistically significant ∼40% increase in *pol1-236 pol12- 216* mutant, consistent with enhanced resection (Figures 5C and 5D).

These results indicate that the CST complex acts after resection initiation mediated by the MRX/Sae2 complex to subsequently limit the extent of ssDNA by fill-in synthesis through its interaction with Polα- primase.

### CST’s interaction with Polα-primase controls the balance between intermediate and large deletions

Because of its effect on ssDNA at DSB, we predicted that CST’s interaction with Polα-primase was important for repair pathway choice. First, survival after DSB induction was decreased in the *pol1-236 pol12-216* mutant, to an extent similar to *stn1Δ*, and no additive effect was observed when combined with *stn1Δ*, suggesting that CST and Polα act together in repair (Figures 1C and 6A).

**Figure 6.**
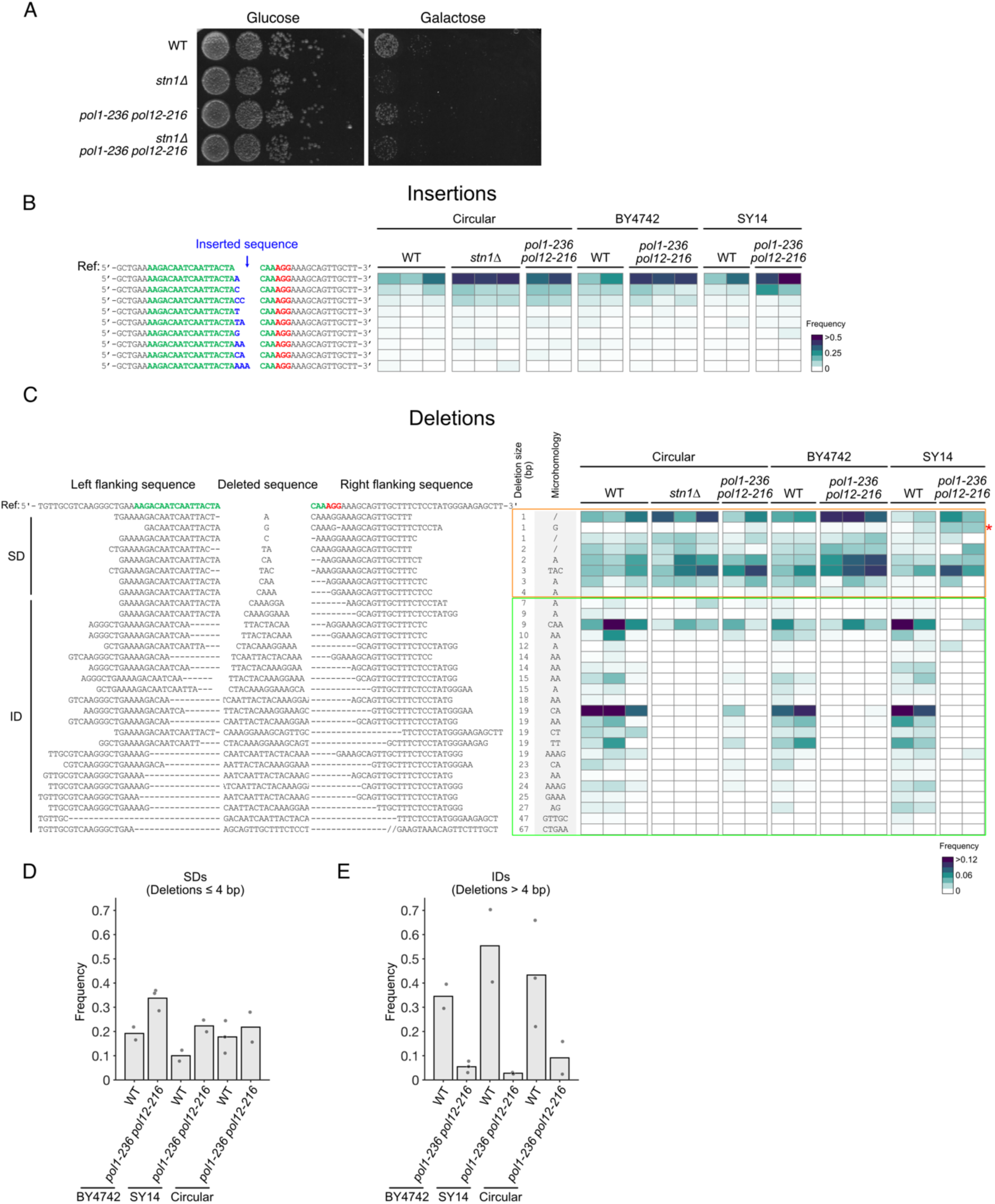
CST’s interaction with Polα-primase is critical for ID/LD balance. (A) Spot assay with the indicated strains showing cell survival and growth with (galactose-containing plate, right) or without (glucose-containing plate, left) Cas9 induction. (B) Heatmap of the frequency of each repair outcome with an insertion for individual experiments (in columns) with the indicated strains, as in Figure 2B. Strains include the parental BY4742 with 16 chromosomes, SY14 with a single linear chromosome, and the strain with a single chromosome, either WT or bearing the *pol1-236 pol12-216* mutation. See Supplemental Data 1 and 2 for the unfiltered data. (C) Heatmap of the frequency of each repair outcome with a deletion for individual experiments with the indicated strains, as in Figure 2C. See Supplemental Data 1 and 2 for the unfiltered data. (D) Frequency of SDs for the indicated strains. Each dot represents an independent experiment. (E) Frequency of IDs for the indicated strains.

We next tested whether CST’s opposite effects on IDs and LDs involved Polα-primase. As observed for *stn1Δ* mutants, the multiplex PCR assay followed by junction mapping revealed that in 4 out of 87 (4.6%) *pol1-236 pol12-216* survivor colonies, unproductive PCRs were due to LDs of 7551 bp mediated by a 22-bp microhomology (Figures 4A and S5).

Using high throughput sequencing of the repaired cut site in survivors, we found that the mutation signature of the *pol1-236 pol12-216* mutant closely resembled that of *stn1Δ* and the CST mutants in general (Figures 6B and 6C). More specifically, the distributions of NHEJ-mediated insertions and SDs were not altered in *pol1-236 pol12-216* compared to WT and *stn1Δ* (Figures 6B, 6C and 6D). In contrast, both NHEJ- and MMEJ-mediated IDs were strongly reduced compared to WT and similar to *stn1Δ* (Figures 6C, 6E, 2A and S2B).

To further test the generality of these observations, we used strains with linear genomes (*i.e.* SY14 with a single chromosome and the parental BY4742 with 16 chromosomes), in which *pol1-236 pol12- 216* mutants are viable. The same mutational signature as in the WT with circular chromosome was found in SY14 and BY4742, indicating that DSB repair was not altered in the single circular chromosome strain compared to linear genomes with telomeres (Figures 6B and 6C). In both SY14 and BY4742 backgrounds, upon introducing the *pol1-236 pol12-216* mutation, no major change in the insertions and the SDs was observed compared to WT (Figures 6B, 6C and 6D). Instead, the mutation signatures were characterized by a marked decrease of IDs (Figures 6C and 6E), mimicking the effects of *pol1-236 pol12-216* and *stn1Δ* mutations in the circular strain.

Altogether, our findings indicate that CST and Polα-primase act together in DSB repair to promote both NHEJ- and MMEJ-mediated IDs. In the absence of CST or when its interaction with Polα is disrupted, these IDs can no longer be formed and repair is either impossible, provoking cell death, or will instead be associated with LDs.

## Discussion

In this work, we report that the CST complex and Polα-primase participate in DSB repair by NHEJ and MMEJ by counteracting resection. Collectively, our findings lead to the following mechanistic model (Figure 7). Once the DSB is formed, recruitment of the MRX and Ku complexes allows the ligation of the two ends through NHEJ, leading to error-free repair. Additional processing of the DSB, in particular if the DSB is not blunt or has other more complex structures, followed by ligation creates limited local mutations such as small insertions and SDs, or SNVs. In all these cases, the CST appears not to be involved, because no ssDNA has been exposed yet. However, the MRX-Sae2 complex also initiates resection, which can kinetically compete with NHEJ. The CST complex, together Polα-primase, can then be recruited to the ssDNA, which would initiate fill-in synthesis, limit the extent of ssDNA and, importantly, create a stable structure composed of dsDNA and hybrid RNA-DNA close to the cut site. This structure can then be amenable to NHEJ repair and lead to IDs (∼5-85 bp). We propose that the size of the deletion is determined by the position of the CST complex on the resected DNA with respect to the DSB site and on where Polα-primase initiates RNA primer synthesis.

**Figure 7.**
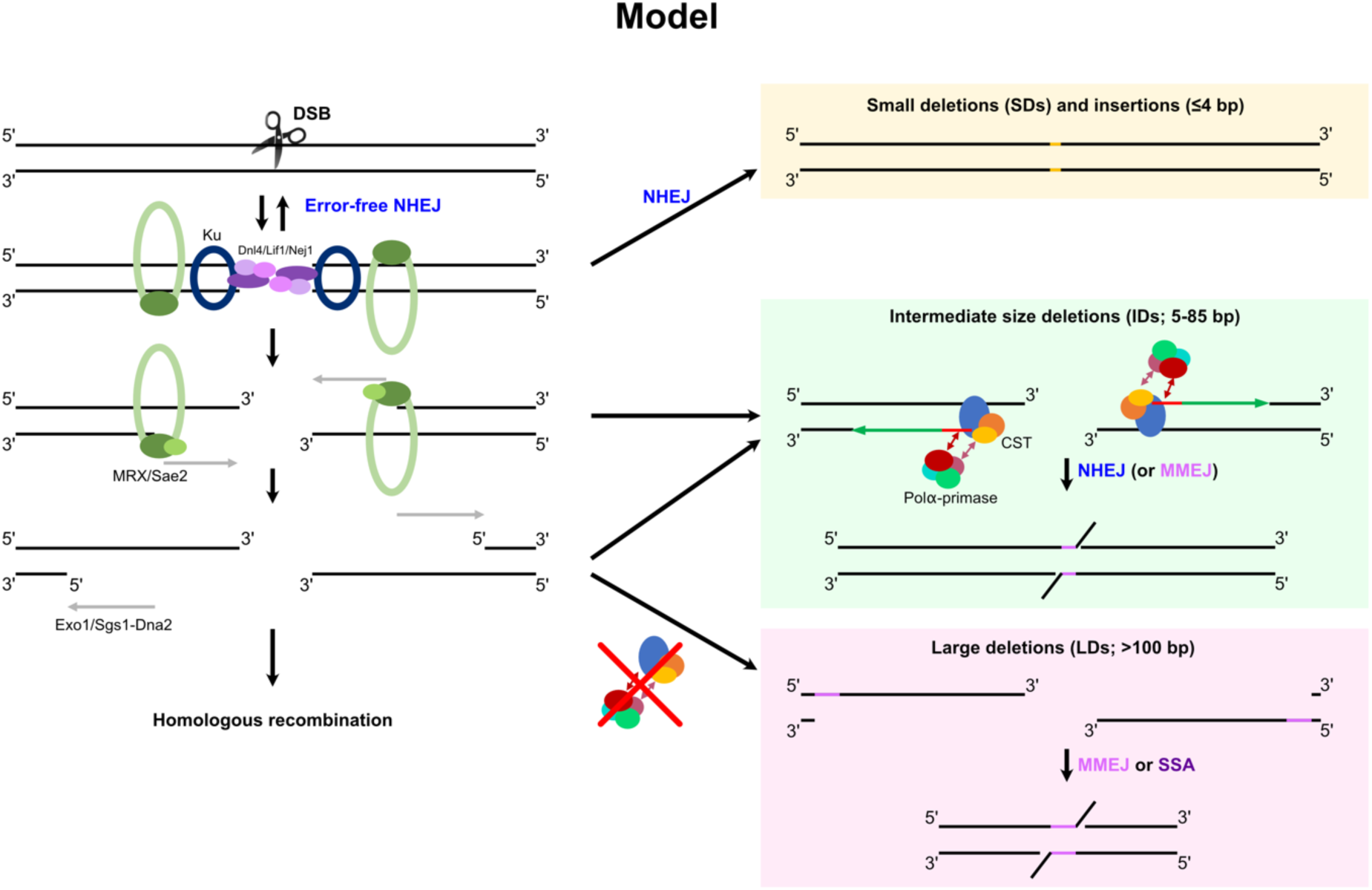
Mechanistic model of CST’s function in DSB repair. The left side shows the canonical repair of a DSB by error-free NHEJ and HR, regulated by the initiation of resection. The right side defines the major mutational events found in this work: (i) before resection initiation, NHEJ can lead to the formation of SDs, small insertions and SNVs; (ii) after resection initiation, the CST complex recruits Polα-primase for fill-in synthesis in a back-up NHEJ pathway or for MMEJ repair, thus leading to IDs; (iii) in the absence of CST or if the interaction between CST and Polα-primase is impaired, extensive resection promotes MMEJ- or SSA-mediated LDs and other rearrangements.

When the CST complex is absent, no double-stranded structure is formed near the break site and resection is not counteracted. NHEJ repair is then no longer available as a repair outcome. Nonetheless, homologies exposed during resection can mediate MMEJ- or SSA-dependent LDs (0.1-20 kb). We assume that other homology-dependent repair mechanisms, *e.g.* HR or break-induced replication, could also be used depending on the sequence context and the presence of homology elsewhere in the genome. In the experimental setting we use where HR is not an available outcome, we also evidence a decrease in cell survival, most prominently in *stn1Δ* and in *pol1-236 pol12-216*, indicating that extensive resection is not always salvaged and becomes toxic. Interestingly, *stn1Δ* displayed a stronger impact than *cdc13Δ* on survival (Figure 1C). We propose that even in the absence of Cdc13, Stn1 and Ten1 can be recruited to the DSB through their OB folds or by an alternative mechanism, albeit with lesser efficiency, which is reminiscent of the ability of Stn1 and Ten1 to associate with and protect telomeres in a Cdc13-independent manner (Ge et al., 2020; Petreaca et al., 2006).

At the genome level, while the CST complex promotes local IDs, it also prevents larger scale deletions, thereby limiting genome instability overall. Interestingly, in mouse cells, Cas9-based genome editing frequently leads to unexpected large deletions of hundreds of bp to several kb, associated with microhomologies (Kosicki et al., 2018; Owens et al., 2019). It would be of great interest to investigate whether in mammalian cells, the CST-dependent pathway modulates the rate of such unintended outcomes to optimize genome editing approaches.

### CST’s role in DSB repair in linear chromosome strains

We reached our conclusions using a unique experimental system designed to uncouple CST’s roles at DSBs from its telomere-specific functions, *i.e.* without the confounding effects of telomere deprotection and without affecting cell survival. That being said, we were able to generalize our results to the context of linear chromosomes by taking advantage of the interaction mutant *pol1-236 pol12- 216*, which is viable in strains with linear chromosomes (BY4742 and SY14). Indeed, the mutational signature obtained in this mutant after DSB induction in the two linear chromosome strains and in the circular chromosome strain was undistinguishable from *stn1Δ*’s. Importantly, these results suggest that the CST complex is able to act at a DSB even in the presence of telomeres, excluding that titration at telomeres would prevent CST from acting at a DSB. We thus conclude that the function of the CST complex uncovered here should also be relevant in the more physiological context of a strain with linear chromosomes.

### CST-dependent post-resection NHEJ leads to deletions of intermediate size

By leveraging an approach that enables the determination at high-resolution of mutation signatures of DSB repair in several genetic contexts (Hussmann et al., 2021; Schimmel et al., 2021), our results evidence IDs generated through NHEJ, which was unexpected since they would require resection initiation. Indeed, binding of Ku to DNA ends protects from resection and conversely, resection displaces Ku (Symington and Gautier, 2011), leading to the view that resection is a commitment step into homology-dependent mechanisms and away from NHEJ. Here, we show that even after resection, NHEJ can still be used for repair in a CST- and Polα-primase-dependent manner and generate ∼5-85 bp deletions. The size of the deletions suggests that they require MRX-Sae2-mediated short-range resection, which was recently mapped at base resolution at discrete positions < 119 bp away from the DSB (Bazzano et al., 2021). Consistently, in the nuclease-dead *mre11-H125N* and in *sae2Δ* mutants, *STN1* deletion no longer affects survival after DSB. In line with these results, the recruitment of Cdc13 to a DSB was previously found to be Mre11-dependent (Oza et al., 2009).

Mechanistically, CST and Polα-primase counteract resection by limiting the extent of ssDNA. We propose that Polα-primase, recruited by CST, creates a double-stranded hybrid structure close to the DSB and facilitates NHEJ through two possible ways. First, the Ku complex requires double-stranded ends, which can include dsDNA with short overhangs and RNA-DNA hybrids (Zahid et al., 2021). The structure formed by Polα-primase activity might thus be directly amenable to Ku binding and NHEJ repair. Second, this structure might delay the extensive degradation of the 3’ ssDNA and the enhanced stability would promote NHEJ (Frank-Vaillant and Marcand, 2002; Zierhut and Diffley, 2008). We also found that NHEJ would often use the base pairing of 2-4 bp (Kramer et al., 1994; Roth and Wilson, 1986), thus facilitating synapsis. Finally, resolution of the local structure would require degradation of the 3’ flap, degradation of the RNA primer and fill-in synthesis of the resulting gap and other remaining ssDNA stretches. Future investigations will be aimed at characterizing these downstream steps to obtain a full picture of how these deletions are formed.

## Material and Methods

### Yeast strains

All yeast strains used in this work are listed in Supplemental Table 1. All are from the BY4742 background. The strain SY14 with a single linear chromosome, SY13 with 2 chromosomes and the parental BY4742 are kind gifts from Prof. Jin-Qiu Zhou and colleagues (Shao et al., 2018). All plasmids used in this work are listed in Supplemental Table 2. Deletion strains were created using standard PCR-based methods (Longtine et al., 1998). Point mutations were generated using Cas9-mediated gene editing (Anand et al., 2017). Strains were grown in rich YPD (yeast extract, peptone, dextrose) or synthetic complete (SC) media.

### Circular chromosome strain

To create the single circular chromosome strain, we used plasmid pJH2970 (gift from Jim Haber (Anand et al., 2017)) containing the Cas9 gene and a site to clone a guide RNA sequence as a base to build plasmid pZX026 in which the sequences to express two guide RNAs were cloned. They were designed to target the subtelomere of chromosome X-R at position 743778 and the subtelomere of chromosome XVI-L at position 17419 (using the initial chromosome numbers and coordinates before they were fused). Transformation of strain SY14 with pZX026 simultaneously with a chimeric double-stranded DNA (Supplemental Table 3) homologous to both subtelomeres allowed the recovery of transformants that have cut the two subtelomeres and recombined them together, thus yielding a strain with a single circular chromosome, yZX168 (Figures S1A-C).

### Spot assay and survival assay

Strains transformed with plasmid pZX013 or pZX010 to target Cas9 to the 5’ UTR of *LYS2* or the 5’ UTR of *URA3*, respectively, were first grown overnight in YPD containing hygromycin (200 µg/mL) at 30°C, then diluted at optical density OD_600 nm_ = 0.7 in YPLG (yeast extract, peptone, 2% lactic acid, 3% glycerol) media and grown for an additional 24h. Cas9 expression was induced by plating the cells on rich solid media (or SC media lacking the appropriate amino-acid) containing 2% galactose or by addition of 2% galactose in the liquid media. As a control, we plated cells from the same culture on 2% glucose-containing plates or added 2% glucose in the liquid media. For spot assays, 10-fold serial dilutions of the liquid culture were performed before depositing 5 µl per spot. For survival assays, after a first estimate of the survival rate for each strain, the appropriate dilution of the culture was plated on galactose-containing plates so as to obtain between 50 and 500 surviving colonies. An additional 100- fold dilution was plated on YPD plates to calculate the rate.

### Resection assays by quantitative PCR

The yeast strains transformed with plasmid pZX013 were grown as for a survival assay and the Cas9 DSB was induced in liquid media for 6h by addition of 2% galactose. The quantitative resection assay was performed as described in (Zierhut and Diffley, 2008). Briefly, genomic DNA was extracted by standard phenol chloroform method and digested by FokI, which cleaves 0.95 kb away from the DSB site but is unable to cleave ssDNA. Using primers flanking the FokI restriction site, qPCR measured the amount of ssDNA relative to the DNA present at each timepoint, by following the formula: %resected = (100/((1+2^ΔC_t_)/2))/f, where ΔC_t_ is the difference in cycles between FokI-digested and undigested samples, and f is the fraction cut by Cas9 determined by qPCR with primers flanking the cut site. All DNA samples were normalized using qPCR primers targeting *ACT1*.

### Plasmid religation assay

The plasmid religation assay was performed as reported in (Wu et al., 2020). Briefly, plasmid pRS426 (Christianson et al., 1992) containing *URA3* as a selection marker was linearized in vitro with EcoRI and the linear form was migrated by gel electrophoresis, excised from the gel and purified. 60 ng of linear or circular plasmid was transformed into the WT circular chromosome strain, *dnl4Δ* mutant and *stn1Δ* mutant, using selective plates lacking uracil. In parallel, to measure plating efficiency, a 6.8 x 10^4^-fold dilution was plated on non-selective YPD media. To calculate the transformation efficiency, the number of colonies on selective media was divided by the number of colonies on YPD multiplied by the dilution factor.

### Southern blot

To verify subtelomere fusion after creating the circular chromosome, a Southern blot was performed as in (Fallet et al., 2014), but the genomic DNA was digested with HindIII and NdeI, and the radiolabelled probe was generated by random priming on a purified PCR fragment overlapping the fusion site (primers used: oT1735 and oT1736, see Supplemental Table 3). The terminal restriction fragment Southern blot, used to detect telomeres, was performed as described in (Coutelier et al., 2018), except that instead of a radioactive probe, an oligonucleotide probe biotinylated at both ends was used (5′-GGGTGTGGGTGTGTGTGGTGGG-3′; Eurofins Genomics) and detected by chemiluminescence. After hybridization of the probe, the membrane was washed 3 x 5 min in wash buffer (58 mM Na_2_HPO_4_, 17 mM NaH_2_PO_4_, 68 mM NaCl, 0.1% SDS). The membrane was next processed for detection with 3 successive incubations (5, 5 and 30 min) in blocking buffer (Thermo Scientific, Nucleic Acid Detection Blocking Buffer) before a 30 min incubation with alkaline phosphatase-conjugated streptavidin (Invitrogen) diluted in blocking buffer (0.4 µg/mL). The membrane was then washed again 3 x 5 min in wash buffer, incubated 2 x 2 min in assay buffer (0.1 M Tris, 0.1 M NaCl pH9.5) and 5 min in CDP-Star substrate (Applied Biosystems) before imaging with a GelDoc system (BioRad).

### High throughput sequencing of mutation signature

Strains transformed with plasmid pZX013 or pZX010 were cultivated and plated on galactose plates as for survival assays, except that more cells were plated so as to obtain hundreds to thousands of surviving colonies. After pooling of the colonies and genomic DNA extraction, a 231-bp amplicon around the Cas9 DSB site was generated using primer containing adapters and multiplexing barcodes for Illumina sequencing (Supplemental Table 3). The libraries were sequenced on an Illumina MiSeq (2 x 250 bp) platform. Analysis of the mutation signature around the Cas9 DSB was performed as described in (Schimmel et al., 2021), using the SIQ software (van Schendel et al., 2022). The resulting analyses are reported in Supplemental Data 2 and the SIQ classification and characterization of mutational events were used to quantify each type of mutation (Supplemental Data 1).

### Hi-C procedure and sequencing

10^7^ cells in 150 mL of YPD media were fixed with 3% formaldehyde for 20 min at 30°C before the reaction was quenched by adding glycine to 0.125 M final concentration for 20 min at room temperature. Hi-C experiments were performed with a Hi-C kit (Arima Genomics) with a double DpnII + HinfI restriction digestion following manufacturer instructions. Samples were purified using AMPure XP beads (Beckman A63882), recovered in 120 µl H_2_O and sonicated using Covaris (∼300 bp) in Covaris microTUBE (Covaris, 520045). Biotinylated DNA was loaded on Dynabeads™ Streptavidin C1 (Fisher Scientific, 10202333). Preparation of the samples for paired-end sequencing on an Illumina NextSeq500 (2×35 bp) was performed using Invitrogen Collibri PS DNA Library Prep Kit for Illumina and following manufacturer instructions. Paired-end sequencing on an Illumina NextSeq500 (2 x 35 bp) was performed.

### Hi-C processing

Reads were aligned and contact maps generated and processed using Hicstuff (https://github.com/koszullab/hicstuff). Briefly, pairs of reads were aligned iteratively and independently using Bowtie2 (Langmead and Salzberg, 2012) in its most sensitive mode against the reference genome CP029160.1 (https://www.ncbi.nlm.nih.gov/nuccore/CP029160; (Shao et al., 2018)). Each uniquely mapped read was assigned to a restriction fragment. Quantification of pairwise contacts between restriction fragments was performed with default parameters: uncuts, loops and circularization events were filtered as described in (Cournac et al., 2012). PCR duplicates (defined as multiple pairs of reads positioned at the exact same position) were discarded. Pairs were binned at 16 kb resolution and contact maps (in mcool format) were generated using OHCA (Serizay et al., 2024).

## Data availability statement

The high-throughput sequencing datasets generated during the current study will be available in a public repository (European Nucleotide Archive).

## Acknowledgements

We are grateful to Jin-Qiu Zhou for providing the SY13, SY14 and BY4742 strains. We thank Teresa Teixeira for discussions, material and reagents, and Jim Haber for plasmids. We thank Stéphane Marcand for his comments on the manuscript. This work benefited from equipment and services from the iGenSeq core facility, at Institut du Cerveau (ICM), supervised by Yannick Marie. We thank the Biomics core sequencing facility of the Institut Pasteur. Research in ZX’s lab was supported by Ville de Paris (Programme Émergence(s)), the Emergence grant of Sorbonne Université, Ligue Contre le Cancer (Subvention Recherche Scientifique 2022), and Fondation ARC pour la recherche sur le cancer (ARCPJA202160003865 and ARCPGA2023110007341_7967). Research in KD’s lab was funded by Fondation ARC pour la recherche sur le cancer (ARCPJA2022070005353), ANR-18-IDEX- 0001, Université Paris Cité IdEx ANR-18-IDEX-0001 and EDF. This work was also supported by the European Research Council under the Horizon 2020 Program (ERC grant agreement 771813) and Agence Nationale pour la Recherche (ANR-22-CE12-0013-01) to RK.

## Author contributions

Investigation: OI, LD, CB, LM and ZX. Formal analysis: OI, LD, LM, RK and ZX. Conceptualization: KD and ZX. Supervision: RK, KD and ZX. Resources, RK, KD and ZX. Writing – original draft: OI, KD and ZX. Writing – review and editing: all authors.

