## Supplemental Information for "The CST complex mediates a post-resection non-homologous end-joining repair pathway and promotes local deletions"

**Short title:** Role of the CST complex in DSB repair

**Authors:** Oana Ilioaia[1], Liébaut Dudragne[1], Clémentine Brocas[2], Léa Meneu[3,4], Romain Koszul[3], Karine Dubrana[2] & Zhou Xu[1,*]

**Affiliations:**

[1]Sorbonne Université, CNRS, UMR7238, Institut de Biologie Paris-Seine, Laboratory of Computational and Quantitative Biology, 75005 Paris, France.

[2]Université Paris Cité, Inserm, CEA, Stabilité Génétique Cellules Souches et Radiations, F-92260 Fontenay-aux-Roses, France.

[3]Institut Pasteur, CNRS UMR3525, Université Paris Cité, Unité Régulation Spatiale des Génomes, 75015 Paris, France.

[4]Sorbonne Université, Collège Doctoral.

**Correspondence:**

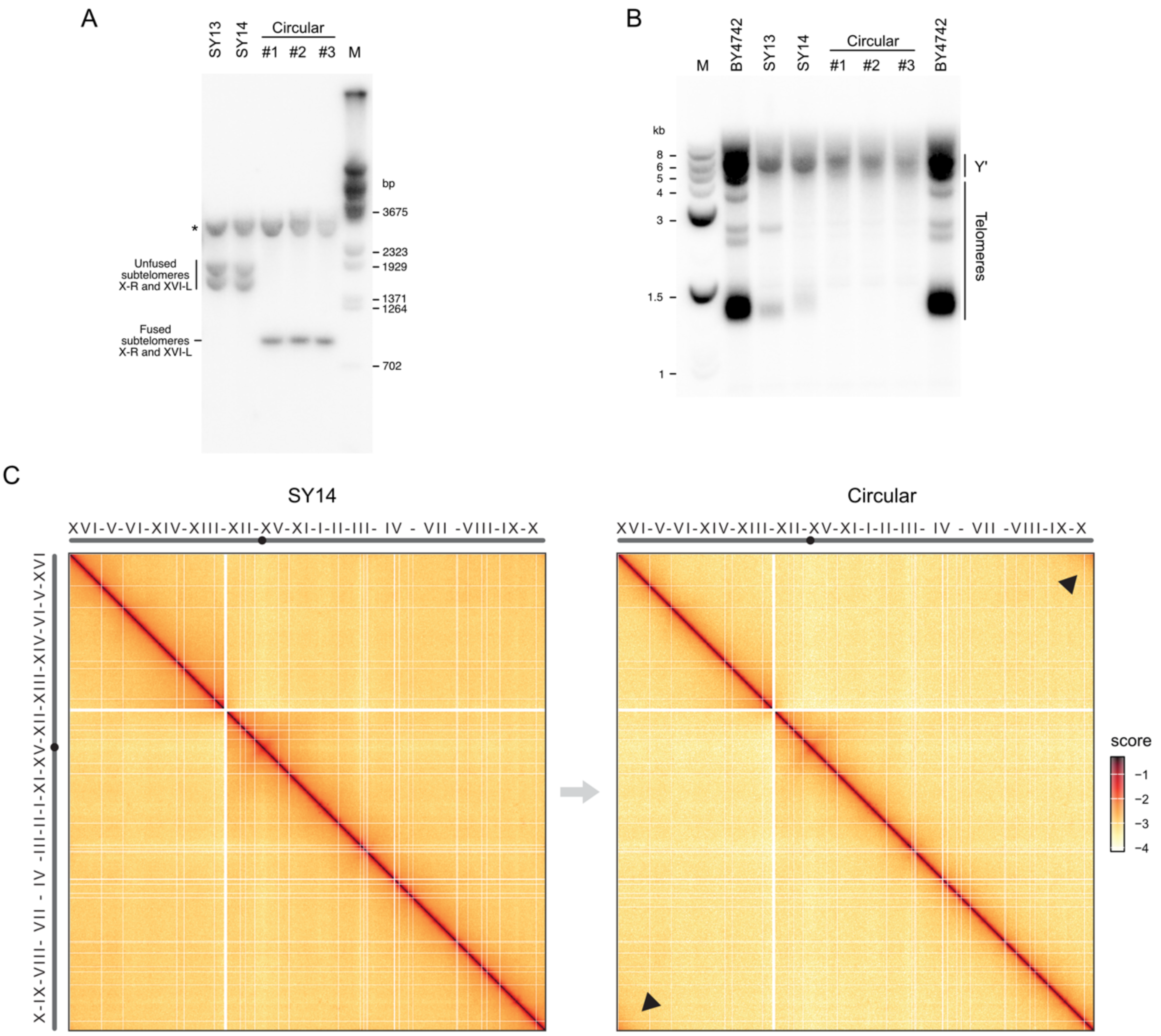


**Supplemental Figure S1. Single chromosome circularization.**

(A) Southern blot for detecting the fusion between the two subtelomeres of SY14. HindIII- and NdeI-digested genomic DNA of the indicated strains was migrated, transferred and probed with a chimeric radiolabeled oligonucleotide complementary to both subtelomeres. Three independent cultures of the circular chromosome strain were tested. M: molecular weight marker ($\lambda$ DNA, BstEII digest). *: non-specific band.

(B) Terminal restriction fragment Southern blot. XhoI-digested genomic DNA of the indicated strains was migrated, transferred and probed with a telomeric probe. M: molecular weight marker.

(C) Normalized Hi-C contact maps of the linear strain SY14 (left) and the single circular chromosome strain (right) with 16-kb resolution. Low to high interaction frequencies are depicted by a color spectrum from light yellow to red. Arrows indicate contacts associated with chromosome circularization.

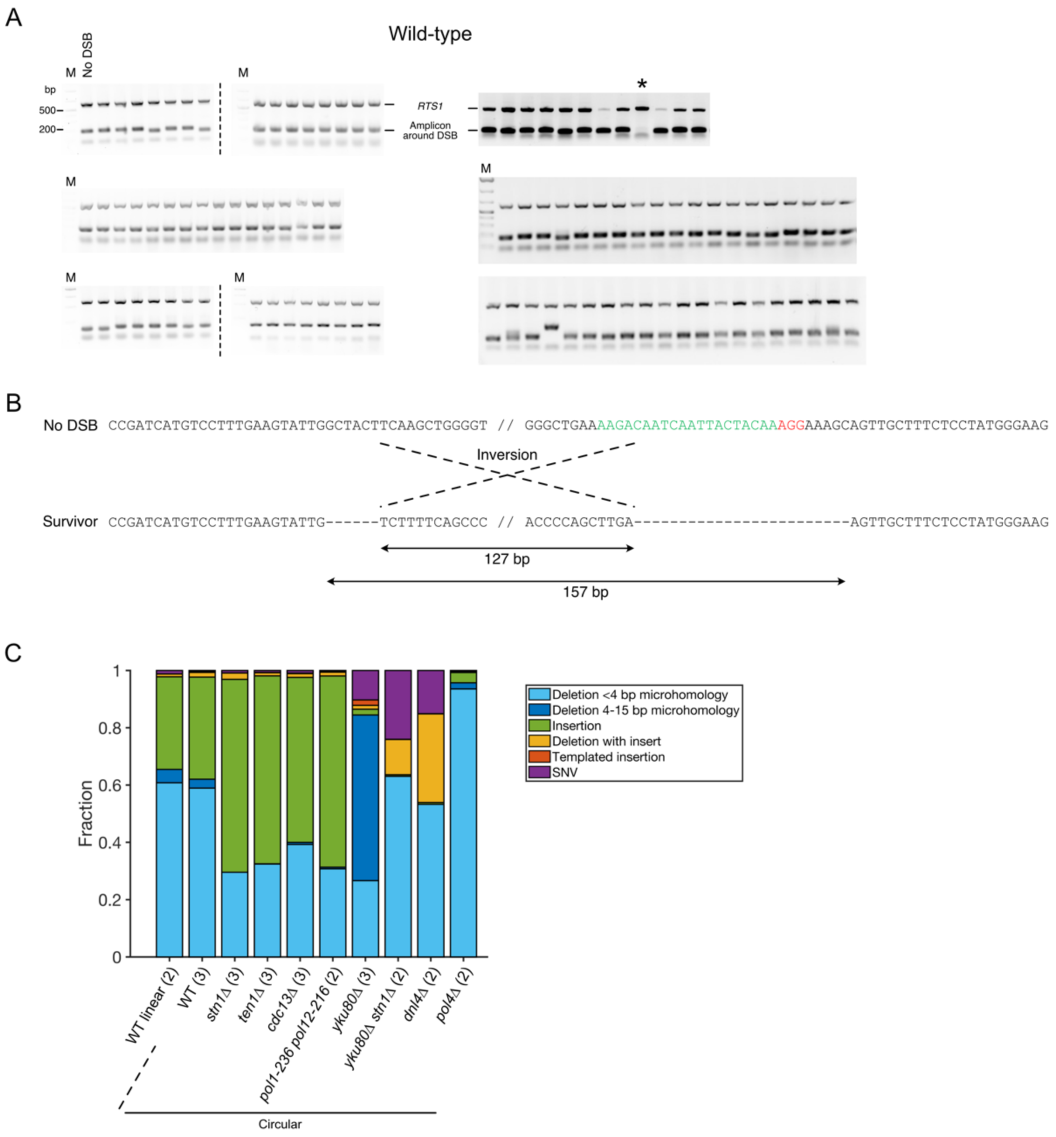


**Supplemental Figure S2. Sequencing of an amplicon around DSB captures nearly all repair event.**

(A) Multiplex PCR showing a 176-bp fragment around the DSB ("Amplicon around DSB") and a fragment in *RTS1*. *: unproductive PCR around the DSB. M: molecular weight marker.

(B) Schematic representation of the single event not captured by PCR in (A). PCR mapping and sequencing of the junction revealed an inversion.

(C) Mutation signature of the indicated strains as shown in Figure 2A, but represented in fraction of each type, without normalization by survival rate.

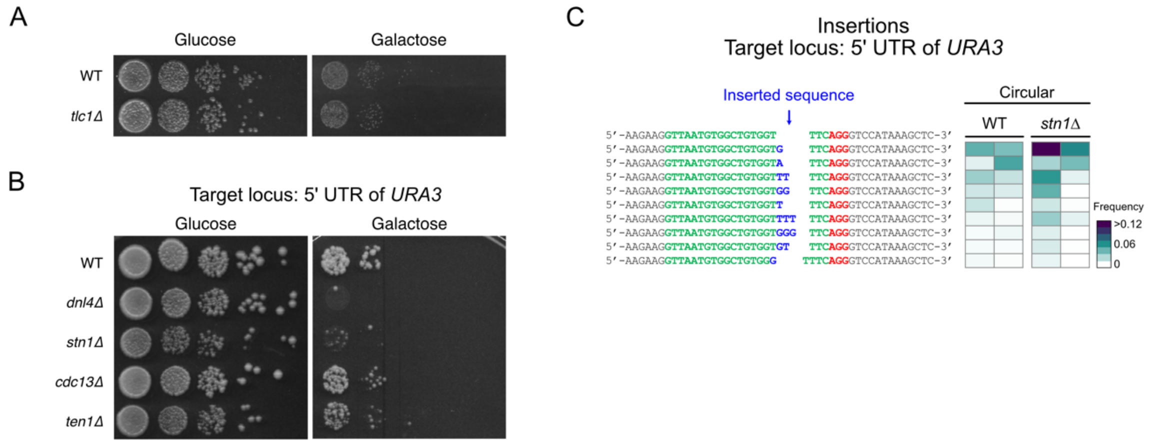


**Supplemental Figure S3. CST's role is telomerase-independent and verified at another DSB locus.**

(A) Spot assay as in Figure 1D with the indicated strains.

(B) Spot assay as in Figure 1D with the indicated strains and with Cas9 targeted to the 5' UTR of *URA3* with a specific guide RNA.

(C) Heatmap of the frequency of each insertion outcome for Cas9 targeted at the 5' UTR of *URA3*, for individual experiments in WT and *stn1Δ* strains. Representation as in Figure 2B, except that only events with frequency > 0.002 are shown. See Supplemental Data 1 and 2 for the unfiltered data.

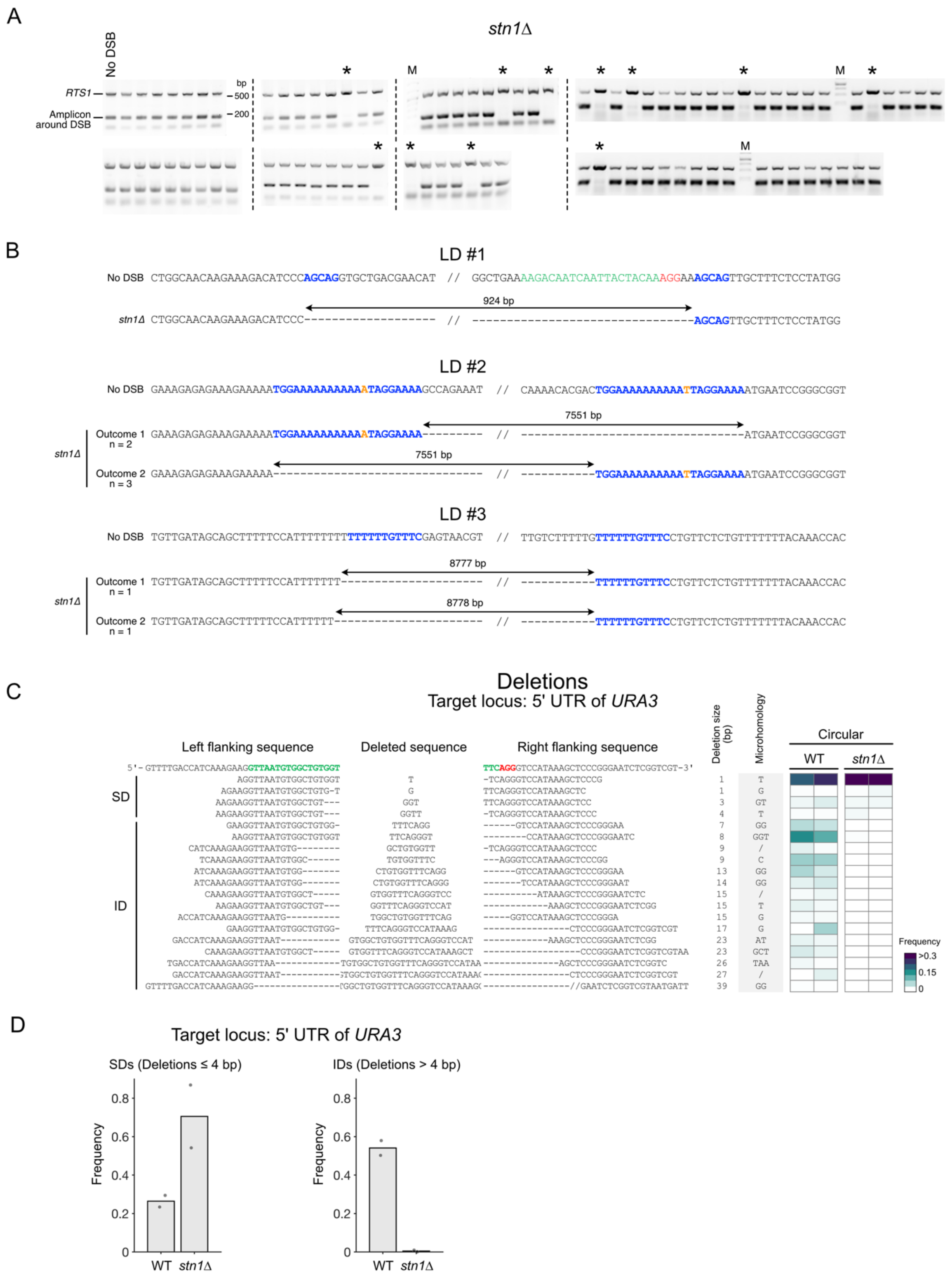


**Supplemental Figure S4. Mutation signature analysis at the DSB targeted to the 5' UTR of *URA3*.**

(A) Multiplex PCR in *stn1Δ* strain showing a 176-bp fragment around the DSB ("Amplicon around DSB") and a fragment in *RTS1*. M: molecular weight marker. * indicate unproductive PCRs around the cut site.

(B) Sequences at the junctions of 8 large deletions detected in (A), revealing LDs of 4 different sizes. The microhomologies used are shown in blue. For LD #2, the microhomology is 22-bp long with a mismatch (in orange). Mismatch repair would eventually resolve the mispairing, leading to the 2 observed outcomes. For LD #3, additional deletion of 1 or 2 bp leads to the 2 observed outcomes.

(C) Heatmap of the frequency of each repair outcome with a deletion for individual experiments (in columns) with the indicated strains, for Cas9 targeted at the 5'UTR of *URA3*. Only deletions that appear with a frequency of > 0.018 in at least one experiment are shown. See Supplemental Data 1 and 2 for the unfiltered data.

(D) Frequency of SDs (left) and IDs (right) for WT and *stn1Δ*, derived from (C), when the DSB is induced at the 5'UTR of *URA3*. Each dot represents an independent experiment.

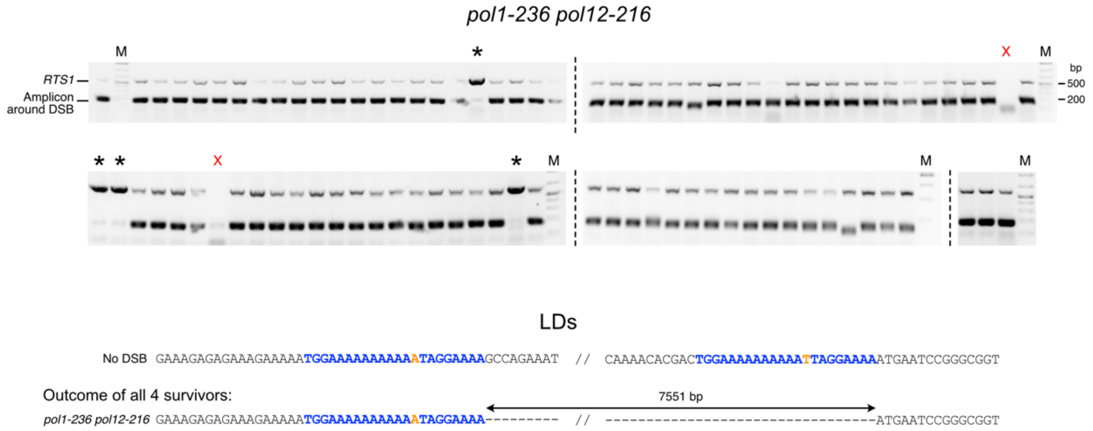


**Supplemental Figure S5. Multiplex PCR assay in *pol1-236 pol12-216* mutant.**

Multiplex PCR in *pol1-236 pol12-216* strain showing a 176-bp fragment around the DSB (“Amplicon around DSB”) and a fragment in *RTS1*. * indicate unproductive PCRs around the DSB. M: molecular weight marker. Red Xs show failed PCR amplification for the control site; the corresponding samples are thus removed from analysis. (Lower part) All 4 unproductive PCRs corresponded to the same LD as in outcome 1 of LD #2 in *stn1Δ* survivor clones, in Figure S4B.

**Supplemental Table 1. List of strains.**

| Strain | Genotype | Figures | Reference |
|---|---|---|---|
| BY4742 | *MATα his3Δ1 leu2Δ0 lys2Δ0 ura3Δ0*; 16 linear chromosomes | S1, 6 | Shao et al., 2018 |
| SY13 | *MATα his3Δ1 leu2Δ0 lys2Δ0 ura3Δ0*; 2 linear chromosomes | S1 | Shao et al., 2018 |
| SY14 | *MATα his3Δ1 leu2Δ0 lys2Δ0 ura3Δ0*; single linear chromosome | 1, S1, 2, S2, 3, 6 | Shao et al., 2018 |
| yZX168 | *MATα his3Δ1 leu2Δ0 lys2Δ0 ura3Δ0*; single circular chromosome | 1, S1, 2, S2, 3, S3, 4, S4, 5, 6 | This work |
| yZX170 | yZX168 *stn1::HIS3* | 1, 2, S2, 3, S3, 4, S4, 5, 6 | This work |
| yZX226 | yZX168 *cdc13::HIS3* | 1, 2, S2, 3, S3, 4 | This work |
| yZX384 | yZX168 *ten1:::HIS3* | 1, 2, S2, 3, S3, 4 | This work |
| yZX207 | yZX168 *dnl4::LEU2* | 1, 2, S2, 3, S3 | This work |
| yZX208 | yZX168 *stn1::HIS3 dnl4::LEU2* | 1, 3 | This work |
| yZX285 | yZX168 *tlc1::HIS3* | 1, S3 | This work |
| yZX283 | yZX168 *pol4::HIS3* | 1, 2, S2 | This work |
| yZX274 | yZX168 *yku80::LEU2* | 1, 2, S2, 5 | This work |
| yZX271 | yZX168 *stn1::HIS3 yku80::LEU2* | 1, 2, S2, 3, 4 | This work |
| yZX292 | yZX168 *pol1-D236N pol12-G325D* | 1, 2, S2, 4, 5, 6, S5 | This work |
| yZX206 | SY14 *dnl4::LEU2* | 3 | This work |
| yZX320 | yZX168 *mre11-H125N* | 5 | This work |
| yZX321 | yZX168 *stn1::HIS3 mre11-H125N* | 5 | This work |
| yZX408 | yZX168 *sae2::LEU2* | 5 | This work |
| yZX415 | yZX168 *stn1::HIS3 sae2::LEU2* | 5 | This work |
| yZX353 | yZX168 *ChrII:460851-460853::LEU2[nt1-559] ChrII:480990-480988::LEU2[nt478-1089] ** | 5 | This work |
| yZX368 | yZX168 *stn1::HIS3 ChrII:460851-460853::LEU2[nt1-559] ChrII:480990-480988::LEU2[nt478-1089] ** | 5 | This work |
| yZX455 | yZX168 *pol1-D236N pol12-G325D ChrII:460851-460853::LEU2[nt1-559] ChrII:480990-480988::LEU2[nt478-1089] ** | 5 | This work |
| yZX421 | yZX168 *stn1::HIS3 pol1-D236N pol12-G325D* | 6 | This work |
| yZX432 | SY14 *pol1-D236N pol12-G325D* | 6 | This work |
| yZX392 | BY4742 *pol1-D236N pol12-G325D* | 6 | This work |

**Supplemental Table 2. List of plasmids.**

| Plasmid | Vector | Selection marker | Insert | Purpose | Reference for vector |
|---|---|---|---|---|---|
| pZX010 | bRA66 | *HPH1* | Guide RNA sequence: GTTAATGTGGCTGTGGTTTC | GAL1-driven Cas9 expression targeting 5' UTR of *URA3* | Anand et al. 2017 |
| pZX013 | bRA66 | *HPH1* | Guide RNA sequence: AAGACAATCAATTACTACAA | GAL1-driven Cas9 expression targeting 5' UTR of *LYS2* | Anand et al. 2017 |
| pZX026 | pJH2970 | *HIS3* | 2 guide RNA sequences: AGCCATAATAGCATCCAGAT and TGAAACGCTGCCGTAAGCAG | Cas9 cut at subtelomeres of ChrX-R and ChrXVI-L for SY14 circularization | Anand et al. 2017 |
| pRS426 | | *URA3* | None | Plasmid religation assay | Christianson et al. 1992 |

**Supplemental Table 3. List of primers.**

| Primer name | Forward/Reverse | Sequence | Target locus | Use |
|---|---|---|---|---|
| oT1721 | F | GGAAAGTTTCCACCAGACGCTAAGTGGTAGC CATAATAGCATCCACTTACGGCAGCGTTTCA CTTTGTTTGGAGAACGGTTGTTAACTTG | Chimera between subtelomeres X-R and XVI-L. | Repair donor sequence for chromosome circularization |
| oT1722 | R | CAAGTTAACAACCGTTCTCCAAACAAAGTGA AACGCTGCCGTAAGTGGATGCTATTATGGCT ACCACTTAGCGTCTGGTGGAAACTTTCC | Chimera between subtelomeres X-R and XVI-L. | Repair donor sequence for chromosome circularization |
| oT1735 | F | GCTTATTCTCAAATGGTGAC | XVI-L subtelomere (17.5 kb from end) | PCR to verify chromosome circularization and generate Southern blot probe |
| oT1736 | R | ACTTCCCAATCATGAGGATC | X-R subtelomere (2.1 kb from end) | PCR to verify chromosome circularization and generate Southern blot probe |
| oZX513 | F | CCACCTTGTCAGATTCGAAGG | Locus 0.95 kb away from Cas9 cut site at 5' UTR of LYS2 | qPCR |
| oZX514 | R | GCATTTACCGAAGTTTACTCCG | Locus 0.95 kb away from Cas9 cut site at 5' UTR of LYS2 | qPCR |
| oT976 | F | CTGGTATGTGTAAAGCCGGT | ACT1 | qPCR |
| oT977 | R | ACGTAGGAGTCTTTTTGACCCA | ACT1 | qPCR |
| oZX108 | F | CGCAACAGCCATCACAATCTC | RTS1 | Control PCR for deletion mapping |
| oZX099 | R | ATGTTCACCACATGAGCGTA | RTS1 | Control PCR for deletion mapping |
| oZX526 | F | AAGTATGCTCATCAATCGTTCGG | Flanking Cas9 cut site at 5' UTR of LYS2 | Large deletion mapping by PCR; qPCR |
| oZX525 | R | CAGACTTAGAAAGCTCTTCCCATA | Flanking Cas9 cut site at 5' UTR of LYS2 | Large deletion mapping by PCR; qPCR |
| oZX641 | F | CAAAGTGGTGATAGAGTTCA | Cas9 cut site at 5' UTR of LYS2 -652 bp | Large deletion mapping by PCR |
| oZX642 | R | ACTGTAAATCAGCTGGCGTT | Cas9 cut site at 5' UTR of LYS2 -652 bp | Large deletion mapping by PCR |
| oZX664 | F | TCAGATCGGATGTGCTTTA | Cas9 cut site at 5' UTR of LYS2 -3421 bp | Large deletion mapping by PCR |
| oZX665 | R | GAGTGCTGTAAGGATTGT | Cas9 cut site at 5' UTR of LYS2 -3421 bp | Large deletion mapping by PCR |
| oZX666 | F | TGCAGCTCTTTGGAACATG | Cas9 cut site at 5' UTR of LYS2 -4626 bp | Large deletion mapping by PCR |
| oZX667 | R | ACTTGGCTCTCCATTGCTT | Cas9 cut site at 5' UTR of LYS2 -4626 bp | Large deletion mapping by PCR |
| oZX649 | F | CCGTTCGACAGAACAAACC | Cas9 cut site at 5' UTR of LYS2 +1982 bp | Large deletion mapping by PCR |
| oZX650 | R | GCACAGTTCTCCGACATT | Cas9 cut site at 5' UTR of LYS2 +1982 bp | Large deletion mapping by PCR |
| oZX639 | F | TTCGACACTCCTTATTCAGGAC | Cas9 cut site at 5' UTR of LYS2 +2196 bp | Large deletion mapping by PCR |
| oZX640 | R | AACGTCATTGCTCGGACATGT | Cas9 cut site at 5' UTR of LYS2 +2196 bp | Large deletion mapping by PCR |
| oZX651 | F | CATGGGTAAAGAGAAGTCT | Cas9 cut site at 5' UTR of LYS2 +3235 bp | Large deletion mapping by PCR |
| oZX652 | R | CTTCCACAAAGCAATATCGAT | Cas9 cut site at 5' UTR of LYS2 +3235 bp | Large deletion mapping by PCR |
| oZX425 | F | AATGATACGGCGACCACCGAGATCTACACAC ACTCTTTCCCTACACGACGCTCTTCCGATCT TGCTCATCAATCGTTCGGAC | Flanking Cas9 cut site at 5' UTR of LYS2 | P5 primer for Illumina sequencing |
| oZX426-oZX435 | R | CAAGCAGAAGACGGCATACGAGAT**[INDEX]** GTGACTGGAGTTCAGACGTGTGCTCTTCCGA TCTTTCAGGCAGCAAGTGACCAT | Flanking Cas9 cut site at 5' UTR of LYS2 | P7 primers for Illumina sequencing. INDEX indicates multiplexing barcode sequence. |
| oZX671 | F | AATGATACGGCGACCACCGAGATCTACACAC ACTCTTTCCCTACACGACGCTCTTCCGATCT ACCGAAGTTATCTGATGTAG | Flanking Cas9 cut site at 5' UTR of URA3 | P5 primer for Illumina sequencing |
| oZX672-oZX677 | R | CAAGCAGAAGACGGCATACGAGAT**[INDEX]** GTGACTGGAGTTCAGACGTGTGCTCTTCCGA TCTGCCCCGTAAAATACTGTTAC | Flanking Cas9 cut site at 5' UTR of URA3 | P7 primers for Illumina sequencing. INDEX indicates multiplexing barcode sequence. |

**Supplemental Data 1.** Unfiltered insertions and deletions as shown in Figures 2, 3, 4, and 6, and in Supplemental Figures S3 and S4.

**Supplemental Data 2.** Raw data output from SIQ software for all analyzed strains.
